# MiCBuS: Marker Gene Mining for Unknown Cell Types Using Bulk and Single Cell RNA-Seq Data

**DOI:** 10.64898/2026.03.20.711946

**Authors:** Shanshan Zhang, Yingying Lu, Qianwen Luo, Lingling An

**Affiliations:** Graduate Interdisciplinary Program in Statistics and Data Science, University of Arizona, Tucson, AZ; Department of Biosystems Engineering, University of Arizona, Tucson, AZ; Department of Biostatistics and Epidemiology, University of Arizona, Tucson, AZ

**Author notes:** These authors contributed equally to this work (Shanshan Zhang, Yingying Lu). Corresponding author: Shanshan Zhang, Ph.D.; Lingling An, Ph.D.

**Keywords:** Marker gene, bulk RNA-Seq, incomplete scRNA-seq, unknown cell type, cellular composition

## Abstract

Identifying cell type-specific expressed genes (marker genes) is essential for understanding the roles and interactions of cell populations within tissues. To achieve this, the traditional differential analysis approaches are often applied to individual cell-type bulk RNA-seq and single-cell RNA-seq data. However, real-world datasets often pose challenges, such as heterogeneous bulk RNA-seq and incomplete scRNA-seq. Heterogeneous bulk RNA-seq amalgamates gene expression profiles from multiple cell types and results in low resolution, while incomplete scRNA-seq does not capture some cell types from the tissue, leading to unknown cell types. Traditional methods fail to identify marker genes for such unknown cell types. MiCBuS addresses this limitation by generating Dirichlet-pseudo-bulk RNA-seq based on bulk and incomplete single-cell RNA-seq data. By performing differential analysis of gene expressions on bulk and Dirichlet-pseudo-bulk RNA-seq samples, MiCBuS can identify the marker genes of unknown cell types, enabling the identification and characterization of these elusive cellular components. Simulation studies and real data analyses demonstrate that MiCBuS reliably and robustly identifies marker genes specific to unknown cell types, a capability that traditional differential analysis methods cannot achieve.

**Availability and implementation:** MiCBuS is implemented in the R language and freely available at https://github.com/Shanshan-Zhang/MiCBuS.

## 1 Introduction

Tissues are intricate ecosystems comprised of diverse cell types, each contributing uniquely to the functionality and equilibrium of the whole (Almendro, et al., 2013; Altschuler and Wu, 2010; Carter and Zhao, 2021). Identifying cell type-specific expressed genes (referred to as cell type marker genes) is essential for understanding the functional roles and interactions of different cell populations within tissues. Marker genes are uniquely or predominantly expressed in specific cell types, serving as molecular signatures that define these cells (Stark, et al., 2019). Defining the identity, functionality, and physiological states of diverse cell types can offer crucial insights into cellular heterogeneity, developmental processes, disease states, and potential therapeutic targets (Altschuler and Wu, 2010; Stark, et al., 2019). Both bulk RNA sequencing of individual cell types and single-cell RNA sequencing are employed to identify these marker genes via differential gene expression analysis, each offering distinct advantages and limitations.

Bulk RNA-seq serves as a powerful tool for the identification of marker genes from pre-isolated distinct cell populations within tissues or biological samples (Gondane and Itkonen, 2023; Stark, et al., 2019). Coupled with experimental cell isolation technologies (e.g., flow cytometry, immunohistochemistry, and laser capture microdissection), bulk RNA-seq can measure the transcriptome for these purified cell populations. This individual cell-type bulk RNA-seq can be used to identify marker genes that are uniquely or predominantly expressed in specific cell types, and has been widely used in various fields, including immunology, to define markers for different immune cell subsets (Gondane and Itkonen, 2023; Racle, et al., 2017). Traditional statistical methods for differential gene expression analysis in bulk RNA-seq such as DESeq2 (Love, et al., 2014) and edgeR (Robinson, et al., 2010), typically model count data using negative binomial distributions to account for variability and to identify differentially expressed genes with high accuracy. Individual cell-type bulk RNA-seq provides a relatively straightforward way to identify cell type-specifically expressed genes and has been instrumental in studies where obtaining pure cell populations is feasible (Gondane and Itkonen, 2023). However, these traditional cell isolation techniques are limited by their reliance on specific experimental platforms, skilled technicians, and limited scalability.

scRNA-seq is a pioneering approach for identifying cell type specific marker genes (Hao, et al., 2024; Jovic, et al., 2022). By examining the transcriptome of individual cells within a heterogeneous population, scRNA-seq unveils a nuanced landscape of gene expression profiles. Sophisticated computational algorithms and clustering techniques have been developed for scRNA-seq data analysis, allowing for the dissection of intricate cell populations and the identification of marker genes with cell-type-specific expression patterns (Jovic, et al., 2022; Pullin and McCarthy, 2024). However, because of technical limitations, scRNA-seq may fail to efficiently capture certain cell types from the tissue due to their size, fragility, or low abundance. This can result in an incomplete representation of the cellular diversity within a tissue, leading to incomplete scRNA-seq data and the issue of unknown cell types (Butler, et al., 2018; Finak, et al., 2015; Gondane and Itkonen, 2023; Kolodziejczyk, et al., 2015). In such situations, traditional differential analysis of scRNA-seq data can only identify marker genes specific to captured cell types, but not for unknown cell types. Further, the high cost of scRNA-seq and its specialized sampling requirements can be a significant barrier, making it less accessible for some research groups and situations (Gondane and Itkonen, 2023). Consequently, cellular deconvolution methods have been developed to estimate cell type proportions and cell type-specific gene expression profiles from heterogenous bulk RNA-seq data, allowing the identification of differentially expressed genes (Jaakkola and Elo, 2022; Nguyen, et al., 2024). Such computational methods usually require a reference cell type gene expression profile either built from individual cell-type bulk RNA-seq or scRNA-seq data (Nguyen, et al., 2024). However, these references may not include all cell types in the bulk RNA-Seq tissue and lead to unknown cell types, particularly when the reference is from a different tissue or a different study (Avila Cobos, et al., 2020; Lu, et al., 2023). In such situations, they cannot identify marker genes for the unknown cell types in the tissue. To our knowledge, no statistical method has been developed to identify marker genes for unknown cell types in the above cases.

We developed MiCBuS, a novel framework that harnesses both bulk RNA-seq and incomplete scRNA-seq data to identify marker genes specific to unknown cell types. MiCBuS operates by utilizing both bulk and single-cell RNA-seq data as input. It initially estimates cell type proportions in bulk RNA-seq samples through a scRNA-seq reference-based deconvolution algorithm. Subsequently, it generates Dirichlet-pseudo-bulk RNA-seq samples based on these estimated cell type proportions. By performing differential analysis of gene expression on bulk and Dirichlet-pseudo-bulk RNA-seq samples, we aim to uncover marker genes associated with unknown cell types, enabling the identification and characterization of these elusive cellular components.

## 2 Methods

### 2.1 Overview

MiCBuS identifies marker genes of unknown cell types using bulk and single-cell RNA-seq data (Figure 1). Unknown cell types are those present in bulk RNA-seq but not detected in scRNA-seq reference data. This can result from technical limitations or biological differences, such as unmatched sample sources or collection times (Avila Cobos, et al., 2020; Lu, et al., 2023). Unlike traditional methods that rely on either bulk or single-cell data alone (Hao, et al., 2024; Love, et al., 2014), MiCBuS integrates both. It estimates cell type proportions in bulk RNA-seq using a scRNA-seq reference-based deconvolution approach, then generates Dirichlet-based pseudo-bulk samples from these estimates. By comparing the Dirichlet-pseudo-bulk and original bulk data, MiCBuS identifies marker genes specific to the unknown cell types, referred to as psMarkers. To validate the psMarkers, we use marker genes identified from complete scRNA-seq data available in simulation studies.

**Figure 1.**
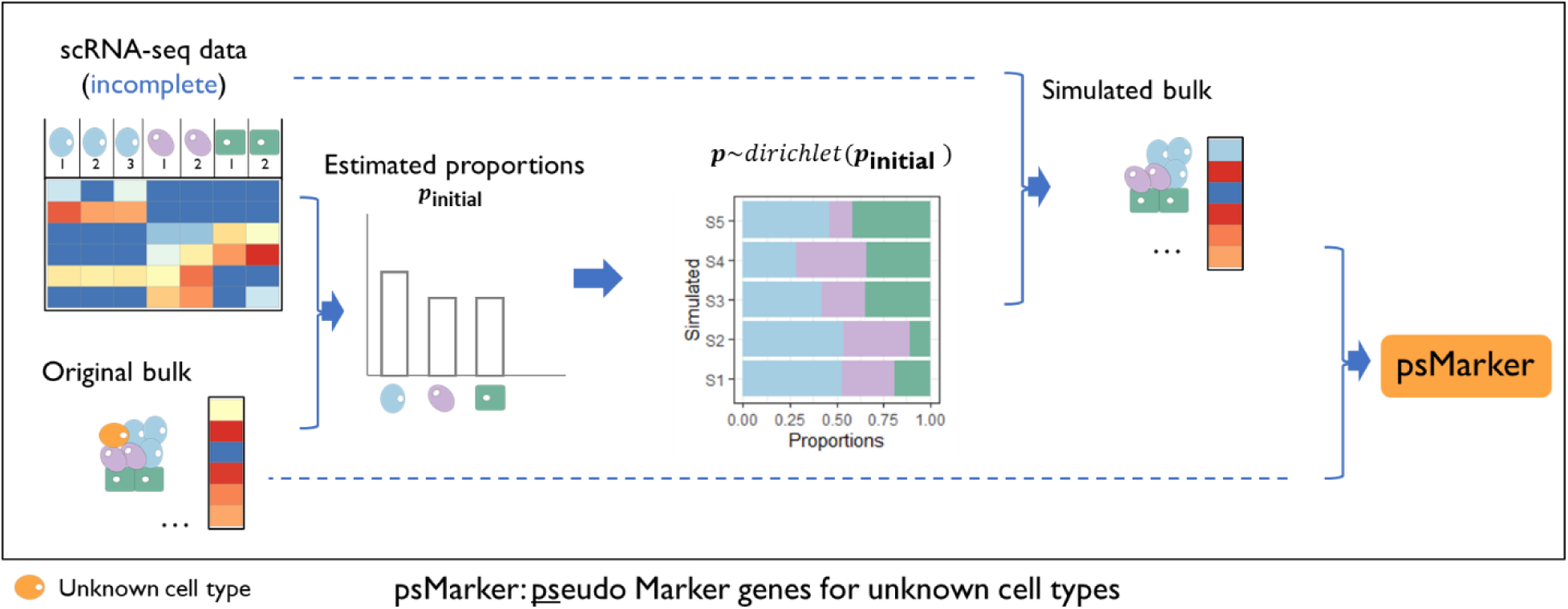
A framework of MiCBuS to identify marker genes for unknown cell types. The unknown cell types, highlighted in orange, are present in bulk RNA-seq data but remain undetected in incomplete scRNA-seq data. MiCBuS takes both bulk and single-cell RNA-seq data as input and follows three major steps. Initially, assuming complete knowledge of all cell types, MiCBuS estimates cell type proportions in bulk RNA-seq data using scRNA-seq as a reference. It then proceeds to generate Dirichlet-pseudo-bulk RNA-seq samples. Notably, during the simulation of cell type proportions ***p***∼*dirichlet*(***p*_initial_** × *s*), the optional parameter of *s* can adjust the spread of the Dirichlet distribution. By comparing these simulated Dirichlet-pseudo-bulk RNA-seq samples with the original bulk RNA-seq data, MiCBuS can identify pseudo marker genes of unknown cell types, referred to as psMarker.

### 2.2 Algorithm

#### 2.2.1 Step 1. Cell type proportion estimation based on scRNA-Seq

When analyzing complete scRNA-seq or bulk RNA-seq data for distinct isolated cell types, traditional methods such as Seurat (Hao, et al., 2024) and DEseq2 (Love, et al., 2014) effectively identify marker genes specific to each cell type. However, real-world datasets often pose challenges—scenarios involving incomplete scRNA-seq and heterogeneous bulk RNA-seq. Incomplete scRNA-seq lacks comprehensive cell type coverage, while heterogeneous bulk RNA-seq combines gene expression profiles from multiple cell types. Employing traditional differential expression analysis techniques on either dataset alone fails to detect marker genes for unknown cell types. If we can use incomplete scRNA-seq to generate random pseudo-bulk RNA-seq data that are comparable to bulk RNA-seq data, except for the exclusion of unknown cell type gene profiles, we can identify genes consistently up-regulated in these simulated datasets. To create such random pseudo-bulk RNA-seq data, we require proportions of known cell types and sufficient variability in cellular compositions across replicated samples.

Although the true cell type proportions of a bulk sample are usually unknown in reality, we can estimate the proportions based on bulk RNA-seq data using a scRNA-seq-reference-based deconvolution approach. In detail, the single-cell RNA-seq data is used as input to generate a single-cell reference matrix from multiple subjects (Lu, et al., 2023). Then, the cell type proportions (referred to as ***p*_initial_**) for bulk RNA-seq data are estimated using a reference-based deconvolution approach. The deconvolution method SECRET with unknown = FALSE is used as a default approach (Lu, et al., 2023), and other reference-based deconvolution methods (Cobos, et al., 2023) can be used as an alternative.

#### 2.2.2 Step 2. Dirichlet-pseudo-bulk RNA-seq data generation

In order to add sufficient variability on cell type proportions for different replicated samples, we assume the proportions are in Dirichlet distribution.

***P****∼dirichlet(α)*

Where the expectation of *p*_*i*_ is

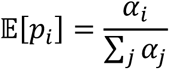

and the variance of *p*_*i*_ is

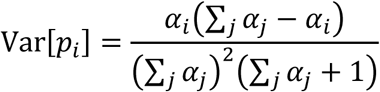

The parameter *α* is defined to be a product of two components (*α* = ***p***_initial_ × *s*), and then the expectation and variance of *p*_*i*_ can be reparametrized as

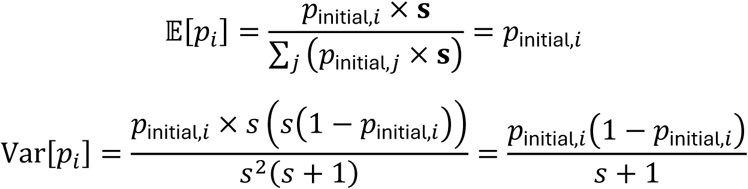

In *α* = ***p***_initial_ × *s*, the first vector ***p*_initial_** is the estimated cell-type proportions calculated from Step 1 and determines the mean of the distribution. Notably, for purpose of ***p*_initial_**estimation, we assume no unknown cell type from Step 1 to Step 2 regardless of the real status of unknown cell types. The second component, *s*, is a parameter to adjust the spread around these mean proportions. A larger *s* results in a more concentrated distribution, indicating less variability around the mean proportions, while a smaller *s* produces a more dispersed distribution, indicating greater variability. When ***p*_initial_** deviates more from the true proportions, a smaller value of *s* can be used to increase the variability to cover the true proportions.

The Dirichlet distribution-based random cell type proportions (***p***_initial_) and the incomplete scRNA-seq data are subsequently utilized in generating Dirichlet-pseudo-bulk RNA-seq data. The generation is processed via the SimBu function of the SimBu R package (Dietrich, et al., 2022). Notably, Dirichlet-pseudo-bulk RNA-seq data differs from original synthetic bulk RNA-seq data in simulation study. The original synthetic bulk RNA-seq data are generated with true cell type proportions in simulation study to mimic real bulk RNA-seq data which is available in real data analysis. Using the original synthetic bulk RNA-seq data in simulation study allows us to have the ground truth for evaluating our proposed method. In contrast, by leveraging the Dirichlet-pseudo-bulk RNA-seq data, we can identify the marker genes for unknown cell types in step 3.

#### 2.2.3 Step 3. Identification of pseudo-marker genes for unknown cell types

The generated Dirichlet-pseudo-bulk RNA-seq data and the original bulk RNA-seq are used as input to detect differentially expressed genes related to unknown cell types via the extensively adopted R package DESeq2 (Love, et al., 2014). The analysis involves setting the contrast between bulk RNA-seq and Dirichlet-pseudo-bulk RNA-seq data, employing default parameters of DESeq2. Specifically, DESeq2 utilizes a negative binomial distribution-based approach to model gene expression counts and applies a Wald test to identify genes exhibiting significant differential expression across varied conditions. The adjusted p-value, FDR, is calculated using Benjamini-Hochberg method. Based on the threshold of FDR and fold change, the genes upregulated in bulk RNA-seq data are taken as pseudo-marker genes for unknown cell types (referred to as psMarker).

### 2.3. Performance evaluation

Since traditional differential analysis approaches lack the capability to identify marker genes for unknown cell types in the above incomplete scRNA-seq scenario, there are no existing methods to compare the performance. Instead, we evaluate the performance of MiCBuS directly based on marker genes in two ways. First, the cellular marker gene annotation database was downloaded from CellMarker2.0, including manually curated cell markers from humans and mice (Hu, et al., 2023). By searching psMarker genes in this database, we assess the reliability of marker gene identification for unknown cell types. Second, we compare psMarker genes with cell-type marker genes using complete scRNA-seq data via traditional gene differential expression analysis methods. This second evaluation is only applicable when the gene profiles of unknown cell types are actually known. Seurat (Hao, et al., 2024) standard procedure is used to do gene differential expression analysis. In detail, we use a nonparametric Wilcoxon Rank Sum test to find markers for every cell type compared to all remaining cells and report only the positive ones (referred to as scMarker). We then use scMarker to evaluate how well the marker genes for unknown cell types are detected in the psMarker gene set. In the performance evaluation, we used the Jaccard index, a commonly used metric to measure how similar two sets of genes are in terms of their shared elements. It is calculated by dividing the size of the intersection of the sets (the number of genes they have in common) by the size of the union of the sets (the total number of unique genes in both sets). The resulting value ranges from 0 to 1, where 0 indicates no overlap, and 1 indicates complete overlap.

## 3 Results

### 3.1 Overview of the proposed method MiCBuS

MiCBuS identifies a marker gene list of unknown cell types using bulk and single-cell (or isolated individual cell-type bulk) RNA-seq data (Figure 1). The unknown cell types refer to the cell types present in the bulk RNA-seq data but not detected by scRNA-seq data. The unknown cell types can be caused by technical limitations of scRNA-seq or biological differences between unmatched samples, as mentioned in Introduction (Avila Cobos, et al., 2020; Lu, et al., 2023). Traditional bulk or single-cell gene differential expression analysis methods fail to identify unknown cell-type specially expressed genes. In this proposed method, MiCBuS takes both bulk and single-cell RNA-seq data as input to study unknown cell types (Figure 1). Through three major steps, including known cell-type proportion estimation, Dirichlet-pseudo-bulk generation, and differential expression analysis, MiCBuS can identify unknown cell-type specially expressed genes, referred to as psMarker. The proposed method was evaluated in the following simulation studies and real data analyses. The major difference between simulation studies and real data analyses was the generation strategy on original bulk RNA-seq data. In simulation studies, the original bulk RNA-seq data were aggregated from single cells in scRNA-seq data with preset mixing proportions of cell types. In contrast, real data analyses used the real bulk RNA-seq data as input. Both simulation studies and real data analyses used real scRNA-seq data as the second input.

### 3.2 Simulation study

We used simulation studies to comprehensively evaluate our proposed method MiCBuS across different aspects. Simulation studies included real scRNA-seq data and synthetic bulk RNA-seq data. Based on different strategies to simulate synthetic bulk RNA-seq data, two different simulation studies were evaluated (Supplementary Note). Simulation study 1 used the scRNA-seq data with masking one or multiple cell types to construct synthetic bulk RNA-seq data, and used the synthetic bulk and the same scRNA-seq data (random sampling 90% cells) as input for MiCBuS analysis. While simulation study 2 used the metastatic tumor scRNA-seq data to construct synthetic metastatic tumor bulk RNA-seq data, and used the synthetic metastatic bulk and primary tumor scRNA-seq data as input for MiCBuS analysis. Because cell types from the primary tumor and metastatic tumor are not fully matching, simulation study 2 naturally contain unknown cell types as in reality.

#### 3.2.1 Evaluation on simulated human pancreas

Simulation study 1 is based on scRNA-seq data from the human pancreas (Baron, et al., 2016). Data of six major cell types (alpha, beta, gamma, delta, acinar, and ductal cells) from three healthy subjects were used. We designed four settings to evaluate the performance of MiCBuS on simulated human pancreas bulk samples under various conditions with different levels of unknown cell type information in the scRNA-seq data (Table S1). Setting 0 served as a negative control with complete scRNA-seq profiles of all six cell types, allowing us to assess MiCBuS under ideal conditions where no cell types were missing. Setting 1 to 3 tested on unknown cell types from one to three with low to high challenging, giving scenarios from simple to extreme missing data condition.

As a result, MiCBuS has strong performance in all four designed settings of simulated human pancreas bulk samples. Setting 0 served as a negative control and did not identify statistically significant differentially expressed genes, indicating that MiCBuS does not generate false positive markers when no cell types are missing (Table S2). Setting 1 tested with one unknown cell type, MiCBuS successfully identified hundreds of upregulated genes as markers for the unknown cell type, validating its effectiveness (Table S2). Setting 2 was designed with two unknown cell types, providing a more challenging condition to test the method’s robustness (Table S2, Figure 2-4, S1). Setting 3, the most extreme scenario, involved three unknown cell types, testing the limits of MiCBuS in identifying marker genes under significant missing data conditions. MiCBuS managed to identify a substantial number of markers even with three missing cell types, demonstrating its robustness (Table S2).

**Figure 2.**
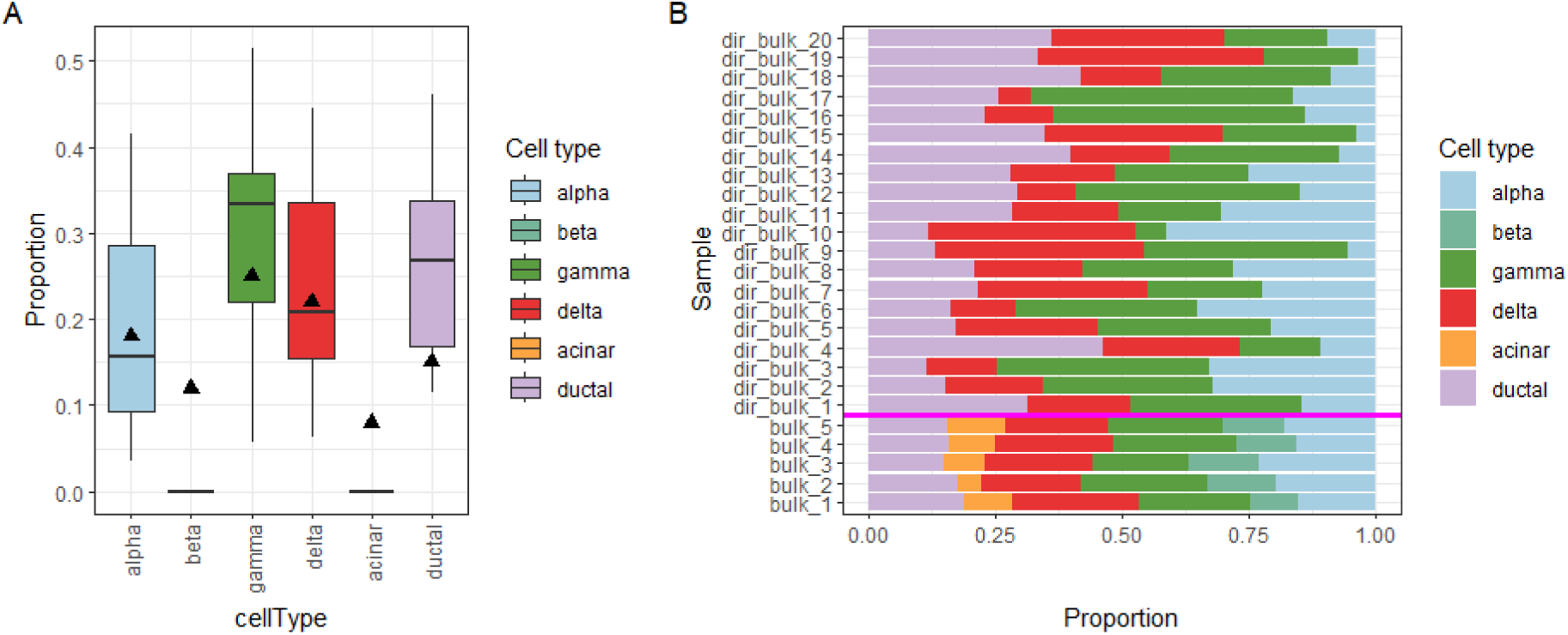
Generation of Dirichlet-pseudo-bulk RNA-seq data in the human pancreas simulation study (setting 2, two unknown cell types). (A) Boxplot of cell type proportions randomly generated from the Dirichlet distribution, used to generate Dirichlet-pseudo-bulk RNA-seq data. The black triangle denotes the true proportion of each cell type, including the unknown cell types (beta and acinar cells). (B) Bar plot of two groups of simulated bulk RNA-seq samples, including 5 synthetic bulk RNA-seq samples, each with 6 cell types (below the bottom of the bright pink line, mimicking real original bulk RNA-seq), and 20 simulated Dirichlet-pseudo-bulk RNA-seq samples, each with 4 cell types (above the bright pink line). Notably, the two cell types in light green and orange are missing from the Dirichlet-pseudo bulk samples.

Using the moderate scenario (setting 2) as illustration, we set the two unknown cell types as beta and acinar cells, which have true proportions of 12% and 8%, respectively (Figure 2A). Following the steps of MiCBuS described in the framework (Figure 1), the initial cell type proportions of original bulk RNA-seq data are estimated using the scRNA-seq reference-based deconvolution approach and then used to generate Dirichlet proportions (detail in Methods). During the generation of random-pseudo-bulk RNA-seq samples, we assumed no unknown cell type. Thus, the Dirichlet-pseudo-bulk RNA-seq samples are only composed of four cell types (Figure 2). In this extreme scenario of two cell types of unknown with large proportion, as expected, the estimated proportions by complete-reference-based cellular deconvolution are inflated (Figure 2A). However, the generated Dirichlet proportions of most cell types are still able to cover the true proportion, including the dramatically affected ductal cells (Figure 2A). Using DEseq2, we identified 1,142 differentially expressed genes between those two groups of bulk RNA-seq samples (FDR < 0.05). We extracted 815 pseudo-bulk upregulated genes as psMarker (log2FoldChanges > 0) for the two unknown cell types and did validation on the top 400 psMarker genes in the following. Using a heatmap on gene expression profiles of complete scRNA-seq data, we found the general expression of psMarker genes is upregulated in beta or acinar cells as compared to the other four cell types (Figure 3A). Searching psMarkers on a public cell type annotation database (Hu, et al., 2023), we found 33 marker genes of the beta cells, such as NKX6.1 (NK6 Homeobox 1) (Figure 3A). NKX6.1, a transcription factor, has been reported to be expressed in beta cells and is essential for beta cell development and function (Marselli, et al., 2010). We also found 63 annotated marker genes for acinar cells (Figure S4). On the other hand, we used the cell-type marker genes identified from complete scRNA-seq data (the top 200 scMarker genes for each cell type) to validate those psMarker genes. These scMarker genes can separate into three categories based on the unknown cell type setting in the incomplete scRNA-seq data: scMarker of beta cells, scMarker of acinar cells, and scMarker of other cell types. We used the Venn diagram to show the tendency of overlapping among those gene sets (Figure 3B). Among the top 400 psMarker genes, 61 genes are in the beta scMarker category with a Jaccard index of 0.113, and 131 genes are in the acinar scMarker with a Jaccard index of 0.279 (Table S2 and Figure 3B). In contrast, none of those psMarker genes are identified in the unique other scMarker category (Table S2 and Figure 3B). Furthermore, psMarker includes 208 more unique genes that are not identified in the above beta and acinar scMarker gene sets (Figure 3B). The heatmap of those genes shows that the expression of a portion of them is still upregulated in beta or acinar cells from complete scRNA-seq data (Figure 3C). To test the stability of MiCBuS, we conducted this simulation 20 times and validated the performance stability of MiCBuS in marker gene identification for the unknown cell type using the Jaccard index as a metric (Figure 4). To test the robustness of our method, we added different levels of noise into the synthetic bulk data in setting 2 to mimic realistic bulk samples. The noise is generated following a normal distribution, as described in Supplementary Notes. After conducting this simulation 20 times, the Jaccard index plot shows that MiCBuS can consistently identify the marker genes for the unknown cell types (Figure 4). This illustrates that our proposed method can properly and robustly identify marker genes for the unknown cell type using bulk and incomplete scRNA-seq data.

**Figure 3.**
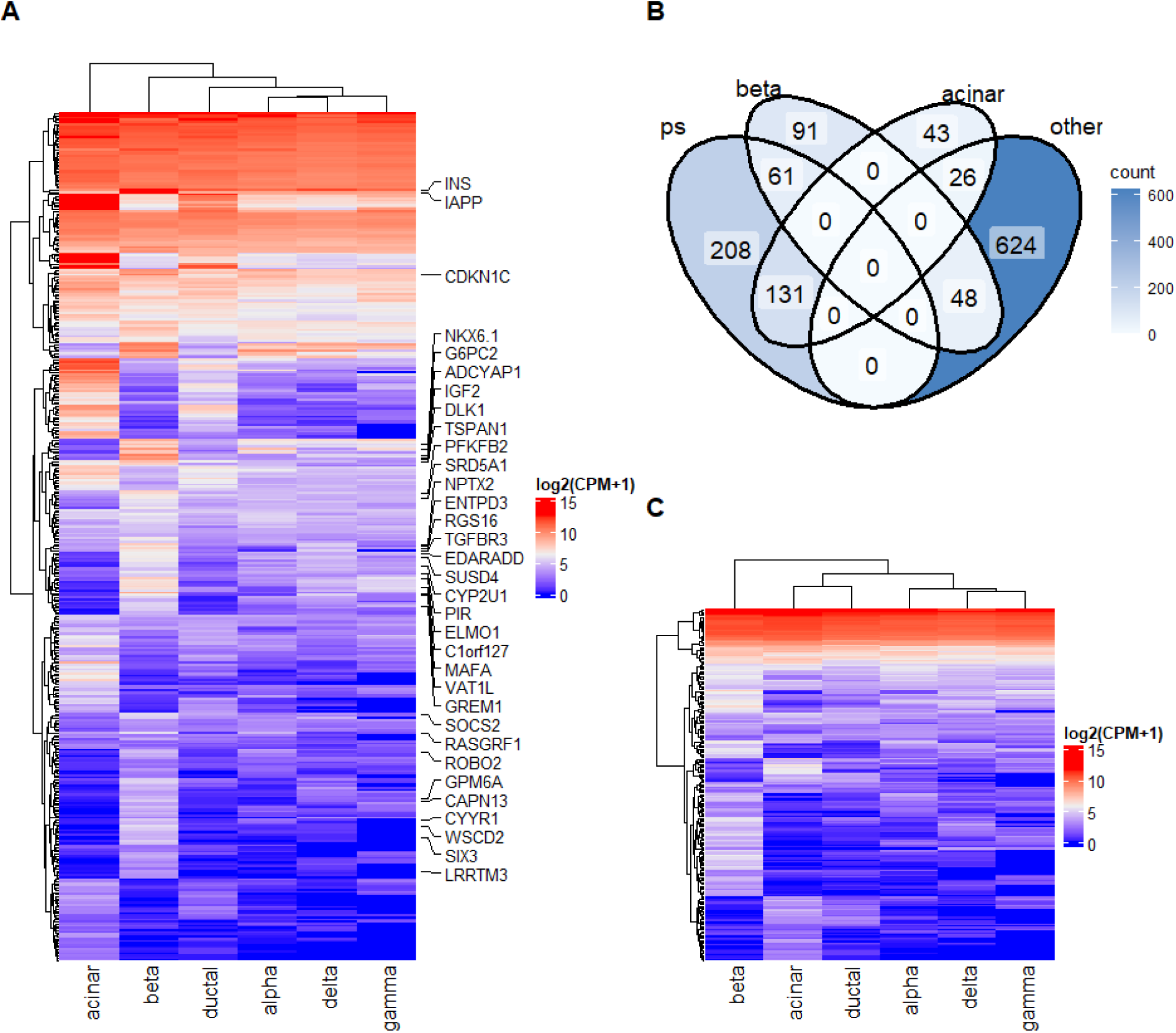
Validation of identified pseudo-marker genes for unknown cell types in the human pancreas simulation study (setting 2, beta and acinar cells). (A) Heatmap of psMarker gene expression using the unmasked (complete) scRNA-seq data. Gene names are shown for those well-known beta cell marker genes based on a public cell type annotation database. In this simulation study, after psMarker gene identification by MiCBuS, scRNA-seq data is unmasked on unknown cell types (beta and acinar cells), providing complete cell type expression profiles for validation purposes. (B) Venn diagram of four gene sets, ps (psMarker genes of unknown cell types identified by MiCBuS) and scMarker (acinar: acinar marker genes; beta: beta marker genes; other: marker genes of the other four known cell types identified using unmasked scRNA-seq data.) (C) Heatmap of psMarker genes (208 unique genes from the Venn diagram in B), showing their expression in the unmasked scRNA-seq data.

**Figure 4.**
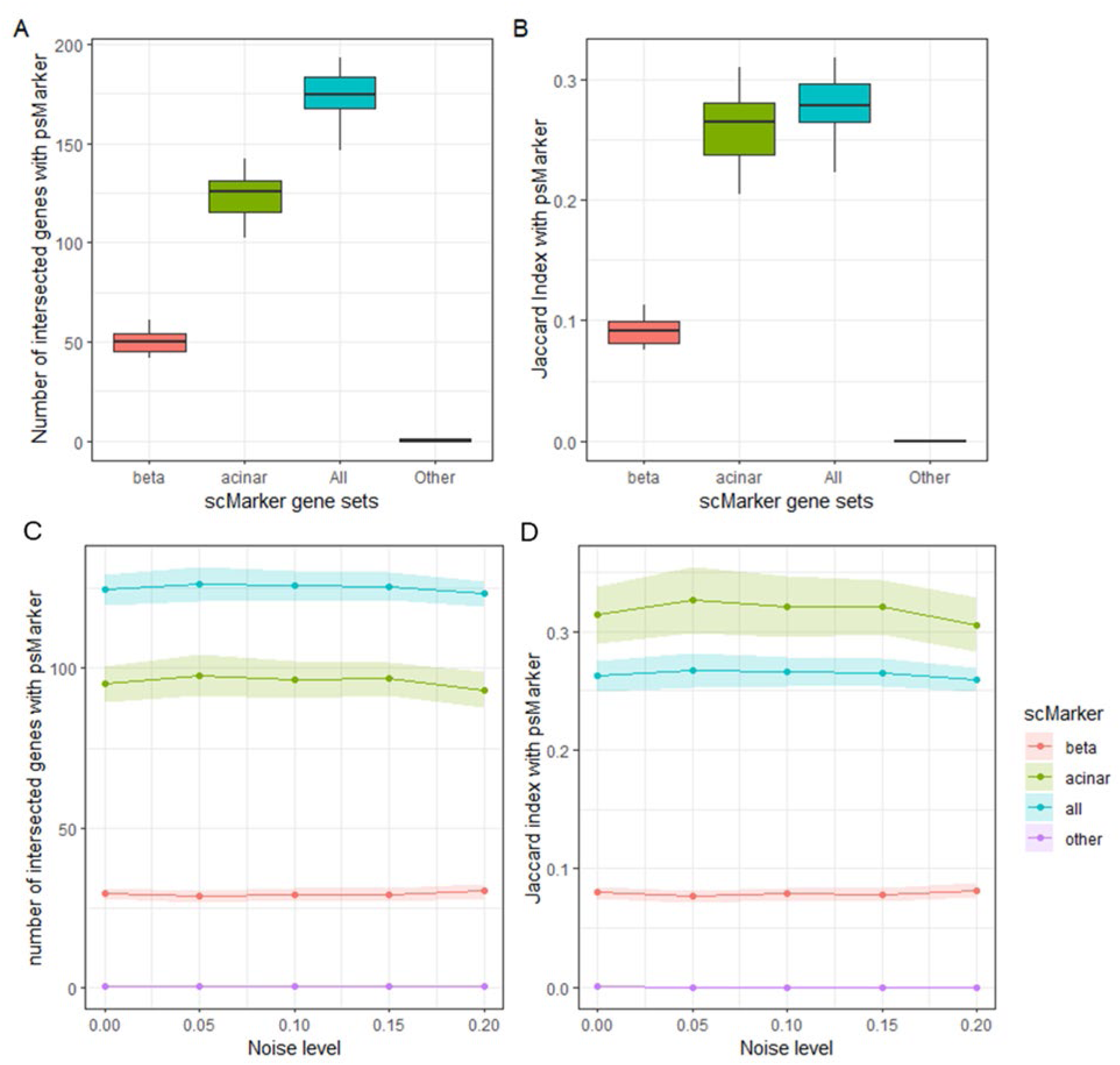
Performance of MiCBuS on unknown cell marker gene identification in 20 runs of the human pancreas simulation study (setting 2, beta and acinar cells). Boxplots of (A) the number of intersected genes with psMarker genes and (B) the corresponding Jaccard index for different subgroup of scMarker genes. Performance of MiCBuS under different levels of noise: (C) the number of intersected genes with the psMarker genes and (D) the corresponding Jaccard index. In all analyses, the beta and acinar cells are the two unknown cell types, and the top 400 psMarker genes are used for evaluation. scMarker gene sets are identified by the unmasked (complete) scRNA-seq data, with the top 200 genes for each cell type are used for evaluation: beta denotes scMarker gene set of beta marker gene; acinar denotes scMarker gene set of acinar marker gene; all denotes the union of the scMarker gene sets for beta and acinar marker genes; and other denotes the marker gene of the other four known cell types, excluding duplicate beta and acinar marker genes.

#### 3.2.2 Evaluation on simulated metastatic lung adenocarcinoma

Simulation study 2 is based on the scRNA-seq data from metastatic lung adenocarcinoma, including both primary and metastatic tumor samples (Kim, et al., 2020). The tumor samples from primary and metastatic sites usually share some common cell types and also often contain distinct ones (Kim, et al., 2020; Lu, et al., 2023). In this case, the primary tumor samples contain 7 cell types, including B lymphocytes, endothelial cells, epithelial cells, Fibroblasts, MAST cells, myeloid cells, and T/NK cells. The tumor samples from the metastatic site brain have one more cell type, oligodendrocytes. In Simulation study 2, scRNA-seq data of metastatic site brain are only used to generate original bulk RNA-seq samples, while scRNA-seq from the primary tumor is actually used as an input in MiCBuS implementation. This setup introduces greater variability in cell type profiling compared to the conditions in simulation study 1. Our results show that MiCBuS can successfully identify marker genes for the unknown cell type, oligodendrocytes, using bulk and incomplete scRNA-seq data (Table S3 and Figure S2-S3).

### 3.3 Real data analysis

In real data analysis, we used real bulk RNA-seq data to evaluate the proposed method. Another input, the incomplete real cell type RNA-seq data, can be the isolated cell type bulk RNA-seq data or scRNA-seq data. We evaluated the proposed method in both cases. The RNA-seq data on cell mixtures from Cobos et. al. 2023, including cell mixture bulk RNA-seq data, individual cell type bulk RNA-seq data, and scRNA-seq data, are perfect real datasets for our method evaluation (Cobos, et al., 2023). The data includes 6 different compositions of cell mixture. All mixtures have 6 different types of cell lines (referred to as cells, to keep consistency with simulation studies), including three breast cancer lines (T47D, BT474, MCF7), monocytes (THP1), lymphocytes (Jurkat), and stem cells (hMSC). Each mixture contained different proportions of cells and was profiled using bulk RNA-seq with 3 replicates. Simultaneously, a portion of mixtures were profiled using scRNA-seq. Additionally, each cell line sample was profiled using bulk RNA-seq with 3 replicates. We masked one or multiple cell types from the latter two datasets to generate the incomplete cell line bulk RNA-seq data and scRNA-seq data. Since the public cell type annotation database (Hu, et al., 2023) does not include marker genes for those six cell lines, we identified both cell line marker genes by cell line bulk RNA-seq data and scRNA-seq data via DEseq2. We then manually curated the top 200 common marker genes for each cell line, sorted by weighted fold change, and added them to the cell type annotation database.

#### 3.3.1 Performance on cell line mixtures with individual cell type bulk RNA-seq as a reference

In real data analysis 1, we evaluated our proposed method MiCBuS with cell line mixture and individual cell type bulk RNA-seq data in 2 settings, one unknown cell type and two unknown cell types. In setting 1, we set the unknown cell type as THP1 cell and masked it from cell line bulk RNA-seq data (referred to as incomplete cell line bulk RNA-seq profiles; Table S4). In setting 2, we set the unknown cell type as THP1 and Jurkat cells and masked them from cell line bulk RNA-seq data (referred to as incomplete cell line bulk RNA-seq profiles; Figure S9). As a result, MiCBuS has good performance in both settings (Figure 5, S3-S5).

**Figure 5.**
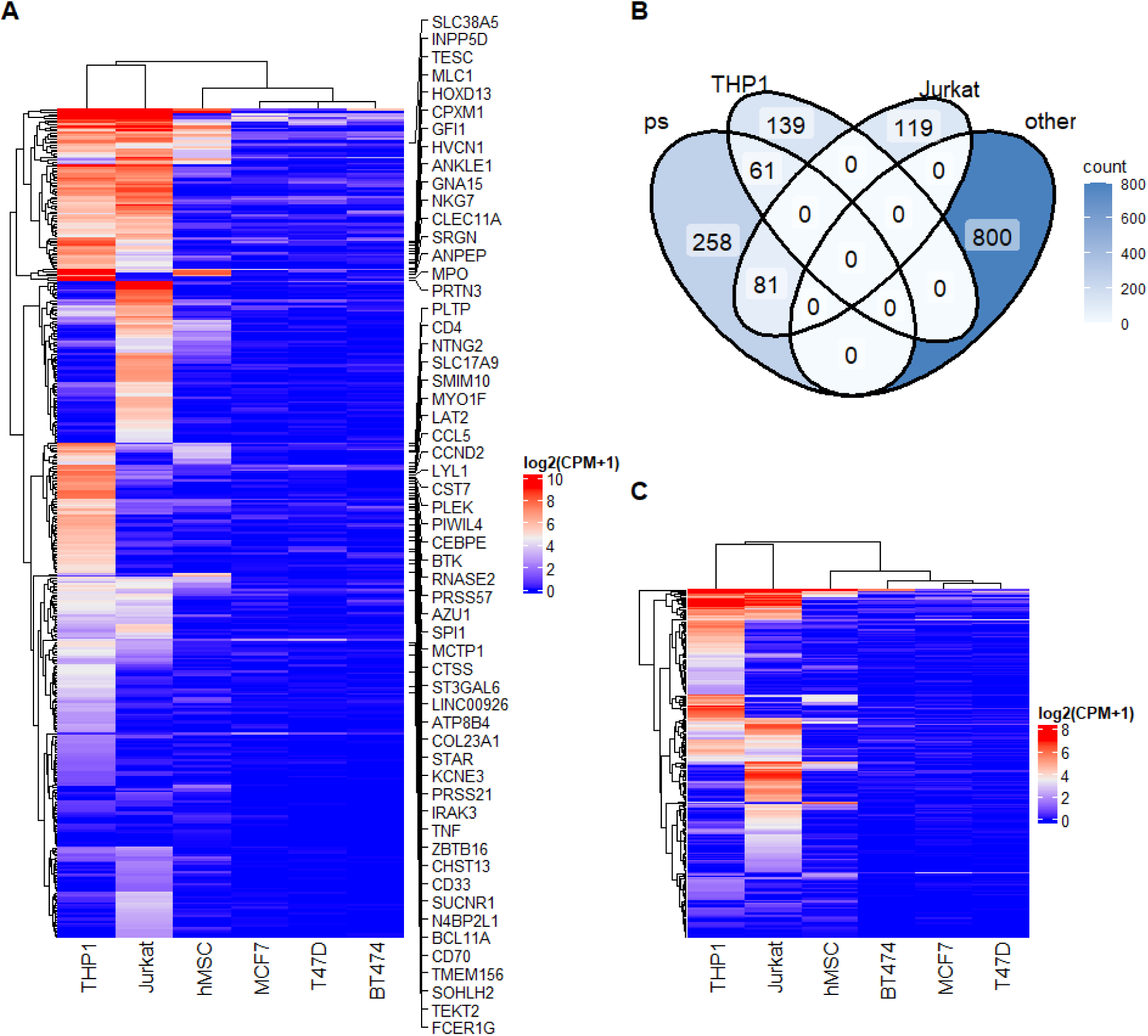
Evaluation of the identified pseudo marker genes in real data analysis 1 (THP1 and Jurkat cells unknown). (A) Heatmap of psMarker gene expression using the unmasked (complete) cell line bulk RNA-seq data. Gene names are shown for genes that are known as marker genes of THP1 cells based on the cell type annotation database. In this setting, the masked unknown cell types are THP1 and Jurkat cells. (B) The Venn diagram of four gene sets, ps (psMarker genes of unknown cell types identified by MiCBuS), and scMarker (THP1: THP1 marker genes; Jurkat: Jurkat marker genes; other: marker genes of the other known cell types identified by complete scRNA-seq data.) (C) Heatmap of psMarker (258 unique genes from Venn diagram in (B)) gene expression using the unmasked cell line bulk RNA-seq data.

We illustrated the real data analysis using the two unknown cell type scenario (setting 2). Following the steps of MiCBuS described in the framework (Figure 1), the steps of MiCBuS on real data analysis are similar to the steps described in simulation study with some minor changes. Under the initial assumption with no unknown cell type, the Dirichlet-pseudo-bulk RNA-seq samples only have four cell types (Figure S3). We generated 10 Dirichlet-pseudo-bulk RNA-seq samples for each type of mixture and then combined them for unknown cell type marker gene mining (Figure S3-S4). Further, we used DEseq2 to identify differentially expressed genes between those two groups of bulk RNA-seq samples. We extracted pseudo-bulk upregulated genes as psMarker (adjusted p-value < 0.05 and log2FoldChanges > 0) for the unknown cell type and did validation on the top 200 psMarker genes in the following. After unknown cell type marker gene mining by MiCBuS, we unmasked THP1 and Jurkat cells from cell line bulk RNA-seq data (referred to as complete cell line bulk RNA-seq data). Using a heatmap on gene expression profiles of complete cell line bulk RNA-seq data, we found the general expression of psMarker genes is upregulated in THP1 or Jurkat cells as compared to the other four cell types (Figure 5A). Searching psMarkers on a manually updated public cell type annotation database (Hu, et al., 2023), we found 57 marker genes of THP1 cells and 66 marker genes of Jurkat cells (Figure 5A and S5). On the other hand, we used the cell-type marker genes identified from complete scRNA-seq data (top 200 scMarker genes for each cell type) to validate those psMarker genes. These scMarker genes can be separated into three categories based on the unknown cell type: scMarker of THP1 cells, scMarker of Jurkat cells, and scMarker of other cell types. We used the Venn diagram to show the tendency of overlapping among those gene sets (Figure 5B). Among the top 200 psMarker genes, 61 genes are in the THP1 scMarker gene set with a Jaccard index of 0.113, and 81 genes are in the Jurkat scMarker gene set with a Jaccard index of 0.156 (Table S4 and Figure 5B). In contrast, none of those psMarker genes are identified in the unique other scMarker category (Table S4 and Figure 5B). Furthermore, psMarker includes 258 more unique genes that are not identified in the above THP1 or Jurkat scMarker gene set (Figure 5B). The heatmap of those genes shows a clear upregulated pattern of gene expression in THP1 or Jurkat cells from complete cell line bulk RNA-seq data (Figure 5C). This illustrates that our proposed method can properly identify marker genes for the unknown cell type using bulk and incomplete cell line bulk RNA-seq data in this real data analysis with two unknown cell types.

#### 3.3.2 Performance on cell line mixtures with scRNA-seq as a reference

In real data analysis 2, we used breast cancer cell mixture bulk RNA-seq data and the corresponding scRNA-seq data to evaluate our proposed method. Compared to cell line bulk RNA-seq data in real data analysis 1, scRNA-seq data introduces more challenges because of large data variability and different sequencing procedures. Although this performance is not as good as real data analysis 1 with cell line bulk RNA-seq data (Table S5, Figure S7-S9), MiCBuS was still able to successfully identify marker genes for the unknown cell type using bulk and incomplete cell scRNA-seq data in this real data analysis 2.

## 4 Discussion

The findings of this study demonstrate that MiCBuS provides a robust solution to the challenge of identifying marker genes for unknown cell types, addressing key limitations of traditional approaches. By integrating bulk RNA-seq and incomplete scRNA-seq data, MiCBuS effectively generates Dirichlet-pseudo-bulk RNA-seq samples to perform differential analysis, allowing for the identification of elusive marker genes that were previously unattainable. This capability is particularly important in scenarios where technical limitations or biological factors result in incomplete scRNA-seq datasets or heterogeneous bulk RNA-seq samples. Simulation studies and real data analyses support the reliability and robustness of MiCBuS in identifying marker genes for unknown cell types, establishing a novel and practical framework for advancing our understanding of cellular heterogeneity.

In simulation studies, synthetic bulk RNA-seq datasets were constructed using either individual cell-type bulk RNA-seq data or scRNA-seq data. The latter posed greater challenges due to its higher variability (Cobos, et al., 2023). For these studies, scRNA-seq datasets from a healthy human pancreas (simulation study 1) and metastatic lung adenocarcinoma (simulation study 2) were used to generate synthetic bulk RNA-seq samples. Various settings, including different numbers of unknown cell types and varying noise levels, were evaluated. Across all settings, MiCBuS reliably identified marker genes for unknown cell types using bulk and incomplete scRNA-seq data. In real data analysis, MiCBuS was further evaluated using more complex real-world datasets. These included bulk RNA-seq data of breast cancer cell mixtures, which were accompanied by cell line bulk RNA-seq data and scRNA-seq data (Cobos, et al., 2023). MiCBuS successfully identified marker genes for unknown cell types using bulk and incomplete cell line bulk RNA-seq data. However, its performance was reduced when using bulk RNA-seq and incomplete scRNA-seq data, likely due to differences in sequencing resolution between bulk and scRNA-seq platforms (Gondane and Itkonen, 2023).

General parameter settings were employed for simulations, with five original bulk RNA-seq samples and 20 Dirichlet-pseudo-bulk RNA-seq samples. The number of bulk RNA-seq samples was chosen to mimic practical scenarios. Results showed that MiCBuS remains effective even with as few as three original bulk RNA-seq samples, as demonstrated in real data analyses. In these analyses, six different mixtures were tested, each containing three replicates. For each mixture, 10 Dirichlet-pseudo-bulk RNA-seq samples were generated and combined to identify marker genes for unknown cell types. MiCBuS successfully identified marker genes across all mixtures, further validating its robustness and adaptability.

The potential applications of marker genes for unknown cell types identified through MiCBuS are vast, particularly in areas where cell type-specific marker genes are critical. For instance, these marker genes can be used in traditional gene differential expression downstream analyses, such as pathway and gene ontology (GO) enrichment, to identify biological processes associated with unknown cell types (Gondane and Itkonen, 2023). Another significant application lies in cellular deconvolution, a computational technique used to infer the cellular composition of bulk RNA-seq data (Avila Cobos, et al., 2020; Jin and Liu, 2021; Sutton, et al., 2022). This method is essential for analyzing complex tissues containing multiple cell types. Previous studies have shown that high-quality marker genes improve the accuracy of deconvolution methods (Momeni, et al., 2023; Nguyen, et al., 2024). Therefore, marker genes for unknown cell types identified by MiCBuS have strong potential for estimating the proportions and gene expression profiles of unknown cell types in heterogeneous bulk RNA-seq samples.

## Declarations

### Ethics approval and consent to participate

Not applicable.

## Consent for publication

Not applicable.

## Availability of data and materials

All datasets used in this study are publicly available. The scRNA-seq data of human pancreas has GEO accession number GSE84133 (Baron, et al., 2016); The scRNA-seq data of human metastatic lung adenocarcinoma has GEO accession number GSE131907 (Kim, et al., 2020); Bulk RNA-seq data of cell mixtures and individual cell lines has GEO accession number GSE220605, and scRNA-seq data of cell mixtures has GEO accession number GSE220606) (Cobos, et al., 2023).

The R code of MiCBuS is available at github.com/ Shanshan-Zhang/MiCBuS.

## Authors’ contributions

SZ and LA conceived the study and designed the methods and algorithms. SZ, LA, YL, and QL contributed to simulation studies. SZ and LA performed the real data analyses. SZ, YL, and LA drafted the manuscript. All authors revised, proofread, and approved the submitted manuscript.

## List of abbreviations

CPM: counts per million; EPIC: a semi-deconvolution method named “Estimate the Proportion of Immune and Cancer cells”; GEO: Gene Expression Omnibus; PREDE: a semi-deconvolution method named “a partial reference-based deconvolution method”; SECRET: a semi-deconvolution method named “Semi-reference based cell type deconvolution”.

## Acknowledgements

We thank Dr. Pavel Sumazin, Mohammad Javad Najaf Panah, and their colleagues (Cobos, et al., 2023) for providing us with additional data relating to their published studies.

## References

Almendro, V., Marusyk, A. and Polyak, K. Cellular heterogeneity and molecular evolution in cancer. Annu Rev Pathol 2013;8:277–302.

Altschuler, S.J. and Wu, L.F. Cellular heterogeneity: do differences make a difference? Cell 2010;141(4):559–563.

Avila Cobos, F., et al. Benchmarking of cell type deconvolution pipelines for transcriptomics data. Nat Commun 2020;11(1):5650.

Baron, M., et al. A Single-Cell Transcriptomic Map of the Human and Mouse Pancreas Reveals Inter- and Intra-cell Population Structure. Cell Syst 2016;3(4):346–360 e344.

Butler, A., et al. Integrating single-cell transcriptomic data across different conditions, technologies, and species. Nat Biotechnol 2018;36(5):411–420.

Carter, B. and Zhao, K. The epigenetic basis of cellular heterogeneity. Nat Rev Genet 2021;22(4):235–250.

Cobos, F.A., et al. Effective methods for bulk RNA-seq deconvolution using scnRNA-seq transcriptomes. Genome Biol 2023;24(1):177.

Dietrich, A., et al. SimBu: bias-aware simulation of bulk RNA-seq data with variable cell-type composition. Bioinformatics 2022;38(Suppl_2):ii141–ii147.

Finak, G., et al. MAST: a flexible statistical framework for assessing transcriptional changes and characterizing heterogeneity in single-cell RNA sequencing data. Genome Biol 2015;16:278.

Gondane, A. and Itkonen, H.M. Revealing the History and Mystery of RNA-Seq. Curr Issues Mol Biol 2023;45(3):1860–1874.

Hao, Y., et al. Dictionary learning for integrative, multimodal and scalable single-cell analysis. Nat Biotechnol 2024;42(2):293–304.

Hu, C., et al. CellMarker 2.0: an updated database of manually curated cell markers in human/mouse and web tools based on scRNA-seq data. Nucleic Acids Res 2023;51(D1):D870–D876.

Jaakkola, M.K. and Elo, L.L. Estimating cell type-specific differential expression using deconvolution. Brief Bioinform 2022;23(1):bbab433.

Jin, H. and Liu, Z. A benchmark for RNA-seq deconvolution analysis under dynamic testing environments. Genome Biol 2021;22(1):102.

Jovic, D., et al. Single-cell RNA sequencing technologies and applications: A brief overview. Clin Transl Med 2022;12(3):e694.

Kim, N., et al. Single-cell RNA sequencing demonstrates the molecular and cellular reprogramming of metastatic lung adenocarcinoma. Nat Commun 2020;11(1):2285.

Kolodziejczyk, A.A., et al. The technology and biology of single-cell RNA sequencing. Mol Cell 2015;58(4):610–620.

Love, M.I., Huber, W. and Anders, S. Moderated estimation of fold change and dispersion for RNA-seq data with DESeq2. Genome Biol 2014;15(12):550.

Lu, Y., Chen, Q.M. and An, L. Semi-reference based cell type deconvolution with application to human metastatic cancers. NAR Genom Bioinform 2023;5(4):lqad109.

Marselli, L., et al. Gene expression profiles of Beta-cell enriched tissue obtained by laser capture microdissection from subjects with type 2 diabetes. PloS one 2010;5(7):e11499.

Momeni, K., et al. Unraveling the complexity: understanding the deconvolutions of RNA-seq data. Translational Medicine Communications 2023;8(1).

Nguyen, H., et al. Fourteen years of cellular deconvolution: methodology, applications, technical evaluation and outstanding challenges. Nucleic Acids Res 2024;52(9):4761–4783.

Pullin, J.M. and McCarthy, D.J. A comparison of marker gene selection methods for single-cell RNA sequencing data. Genome Biol 2024;25(1):56.

Racle, J., et al. Simultaneous enumeration of cancer and immune cell types from bulk tumor gene expression data. Elife 2017;6.

Robinson, M.D., McCarthy, D.J. and Smyth, G.K. edgeR: a Bioconductor package for differential expression analysis of digital gene expression data. Bioinformatics 2010;26(1):139–140.

Stark, R., Grzelak, M. and Hadfield, J. RNA sequencing: the teenage years. Nat Rev Genet 2019;20(11):631–656.

Sutton, G.J., et al. Comprehensive evaluation of deconvolution methods for human brain gene expression. Nat Commun 2022;13(1):1358.

