## Supplemental Information for "MiCBuS: Marker Gene Mining for Unknown Cell Types Using Bulk and Single Cell RNA-Seq Data"

**Supplementary Data**

**Supplementary Notes**

***1 Datasets***

**1.1 Synthetic datasets**

Two simulation studies were conducted to evaluate the proposed method.

- Simulation study 1 was based on scRNA-seq data from the human pancreas (Baron, et al., 2016), which is widely used in deconvolution studies (Avila Cobos, et al., 2020; Dong, et al., 2021). Six major cell types (alpha, beta, gamma, delta, acinar, and ductal cells) were extracted from three healthy individuals. The same scRNA-seq dataset was used to create both the synthetic bulk RNA-seq samples and the reference data (with 90% of cells sampled) for deconvolution.
- Simulation study 2 used scRNA-seq data from metastatic lung adenocarcinoma samples, including both primary and metastatic tumors (Kim, et al., 2020). The tumor samples from primary and metastatic sites usually share some common cell types and very likely have unique cell types as well (Kim, et al., 2020; Lu, et al., 2023). In this case, the primary tumor samples

contain 7 cell types, including B lymphocytes, endothelial cells, epithelial cells, Fibroblasts, MAST cells, myeloid cells, and T/NK cells. The tumor samples from the metastatic site brain have one more cell type, oligodendrocytes. In this setting, synthetic bulk RNA-seq data were generated using cells from the metastatic site, while the reference scRNA-seq data came only from the primary tumor, introducing realistic mismatches between the datasets.

In the silicon generation way, the synthetic bulk RNA-seq was constructed by aggregating single cells from scRNA-seq datasets. In simulation study 1, the simulation strategies are detailed in Table S1. The general parameters across all simulation settings are as follows: alpha = 18%, beta = 12%, gamma = 25%, delta = 22%, acinar = 8%, and ductal = 15%. These values were set solely for method evaluation purposes and do not reflect actual tissue compositions. In simulation study 2, we assigned the unknown cell type oligodendrocytes a 20% proportion, and the remaining cell types reflected realistic proportions inferred from the metastatic brain scRNA-seq dataset: B lymphocytes = 5%, endothelial cells = 1%, epithelial cells = 40%, fibroblasts = 2%, MAST cells = 2%, myeloid cells = 20%, and T/NK cells = 10%.

For each bulk RNA-seq sample, 1,000 cells were pooled. In total, 5 pseudo-bulk RNA-seq samples were generated as the original bulk data, and 20 Dirichlet-pseudo-bulk RNA-seq samples were generated for marker gene identification. To mimic the sampling noise present in bulk RNA-seq, we added random noise from a normal distribution to the true cell type proportions, rescaled them to sum to 1, and used these to construct the original pseudo-bulk RNA-seq data. This process was repeated to generate all pseudo-bulk RNA-seq samples. To simulate scRNA-seq data variability, we randomly sampled 90% of cells from the scRNA-seq dataset. These resampled cells were used for generating Dirichlet-pseudo-bulk RNA-seq samples and identifying marker genes for unknown cell types. Depending on the simulation setting, either complete or partially masked scRNA-seq data (with certain cell types excluded) were used. We drew the proportions $\boldsymbol{p} \sim dirichlet(\boldsymbol{p}_{\text{initial}} \times s)$, where the initial cell type proportions for the bulk samples $\boldsymbol{p}_{\text{initial}}$ were estimated via incomplete scRNA-seq dataset and $s = 10$ was selected as it allows sufficient dispersion around the estimated means.

To test the robustness of our method, different levels of Gaussian noise $N(0, aY)$ were introduced into the synthetic bulk RNA-seq data under simulation setting 1. Here, $a$ was varied across the values $(0,\ 0.1, 0.2, 0.3, 0.4)$, and $Y$ represents the synthetic bulk data.

#### 1.2 Real datasets

In this study, we evaluated our proposed method, MiCBuS, using real bulk RNA-seq data from cell line mixtures, originally generated to benchmark bulk RNA-seq deconvolution methods (Cobos, et al., 2023). These real datasets include six defined cell types: three breast cancer cell lines (T47D, BT474, MCF7), monocytes (THP1), lymphocytes (Jurkat), and human mesenchymal stem cells (hMSC). The bulk RNA-seq data were obtained from six different cell mixtures, each comprising varying proportions of the six cell types and sequenced in triplicates. In parallel, both cell line-specific bulk RNA-seq profiles and single-cell RNA-seq (scRNA-seq) data were generated for the same set of cell lines, enabling reference construction.

To mimic incomplete reference conditions for unknown cell type marker gene identification, we manipulated the available reference datasets in two ways. First, in real data analysis 1, we masked one or two cell types (e.g., THP1 or both THP1 and Jurkat) from the cell line bulk RNA-seq reference. In this setup, the masked data were treated as unknown, and MiCBuS used the remaining reference data to generate Dirichlet-pseudo-bulk RNA-seq samples. These samples were compared to the real bulk RNA-seq data of the cell mixtures to detect pseudo-marker genes (psMarkers) associated with the hidden cell types.

Second, in real data analysis 2, we instead used scRNA-seq data as the reference and introduced incompleteness by masking THP1 cells. This setup represented a more challenging scenario due to the inherent variability and technical differences between scRNA-seq and bulk RNA-seq data. Under this setting, the masked scRNA-seq reference was used to generate Dirichlet-pseudo-bulk RNA-seq samples, and MiCBuS was applied to discover marker genes for the excluded cell type based on expression contrasts with real bulk RNA-seq mixtures.

In both settings, the goal was to identify marker genes specific to cell types missing from the reference data. Validation was conducted using both the complete data (unmasked cell types) and a manually curated marker gene database, assembled from top-ranked differentially expressed genes identified by DESeq2 in both bulk and scRNA-seq profiles.

### Supplementary Tables

**Table S1. Simulation study 1 strategy to create synthetic datasets using real scRNA-seq data from human pancreas.**

| | # of cell types in synthetic bulk | # of cell types in sc data | Marker gene identification |
|---|---|---|---|
| **Setting 0** | 6 | 6 | negative control |
| **Setting 1** | 6 | 5 | one unknown cell type |
| **Setting 2** | 6 | 4 | two unknown cell type |
| **Setting 3** | 6 | 3 | three unknown cell types |

**Table S2. Performance of MiCBuS in simulation study 1**

| Setting | scMarker | Intersected_genes | Jaccard_index |
|---|---|---|---|
| 0 | None | 0 | 0 |
| | Unknown cell type 1 | 115 | 0.404 |
| 1 | Other_unique | 0 | 0 |
| | Unknown cell type 1 | 61 | 0.113 |
| | Unknown cell type 2 | 131 | 0.279 |
| | All unknown | 192 | 0.316 |
| 2 | Other_unique | 0 | 0 |
| | Unknown cell type 1 | 50 | 0.067 |
| | Unknown cell type 2 | 59 | 0.080 |
| | Unknown cell type 3 | 89 | 0.125 |
| | All unknown | 195 | 0.195 |
| 3 | Other_unique | 2 | 0.002 |

Note: Setting 0 did not identify statistically significant differentially expressed genes.

**Table S3. Performance of MiCBuS in simulation study 2**

| scMarker | Intersected_genes | Jaccard_index |
|---|---|---|
| Unknown cell type 1 | 105 | 0.356 |
| Other_unique | 6 | 0.004 |

**Table S4. Performance of MiCBuS in real data analysis 1.**

| Setting | scMarker | Intersected_genes | Jaccard_index |
|---|---|---|---|
| | Unknown cell type 1 | 60 | 0.176 |
| 1 | Other_unique | 2 | 0.002 |
| | Unknown cell type 1 | 61 | 0.113 |
| | Unknown cell type 2 | 81 | 0.156 |
| | All unknown | 142 | 0.216 |
| 2 | Other_unique | 0 | 0.000 |

**Table S5. Performance of MiCBuS in real data analysis 2.**

| scMarker | Intersected_genes | Jaccard_index |
|---|---|---|
| Unknown cell type 1 | 19 | 0.050 |
| Other_unique | 6 | 0.005 |

### Supplementary Figures

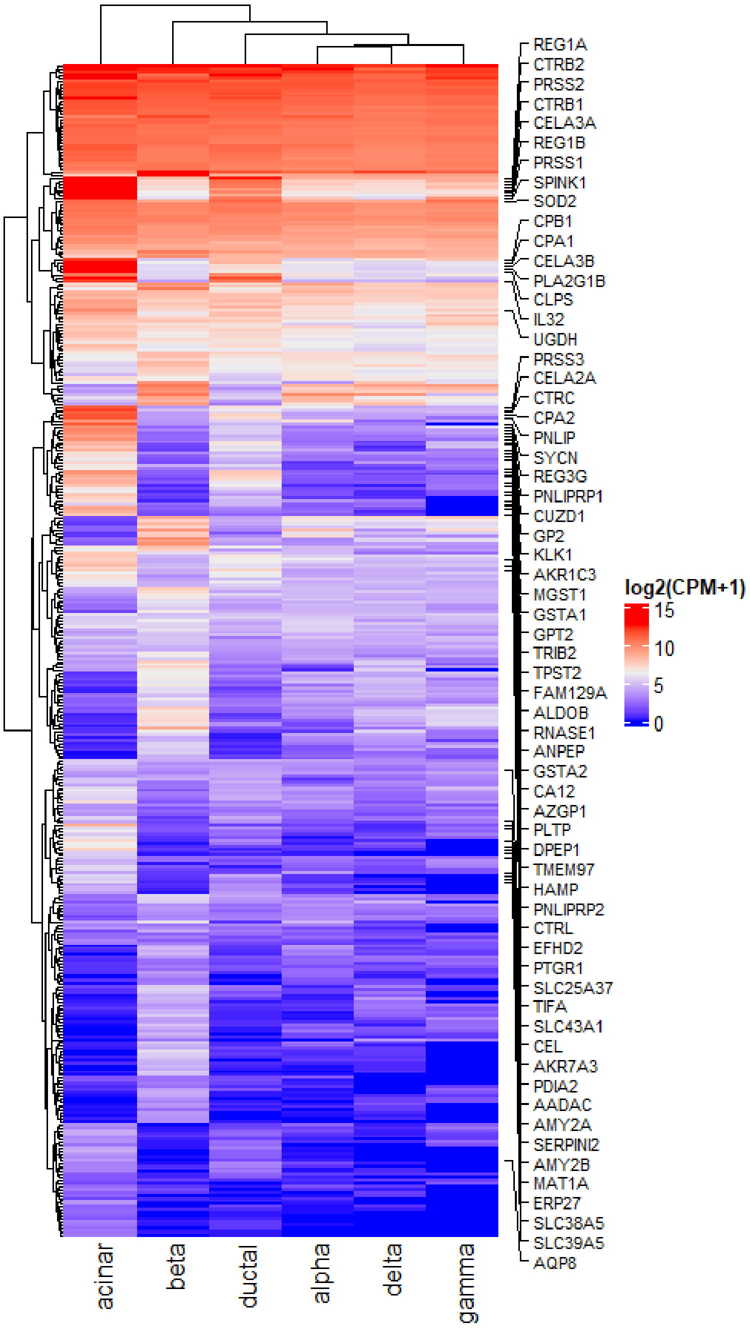


**Figure S1** Identification of the pseudo marker genes in setting 2 for two unknown cell types in simulation study 1. Heatmap of psMarker gene expression using the complete scRNA-seq data. Gene names are shown for genes that are known as marker genes of acinar cells based on the public cell type annotation database. In this simulation study, we know the unknown cell types are beta and acinar cells, and the complete scRNA-seq data is obtainable.

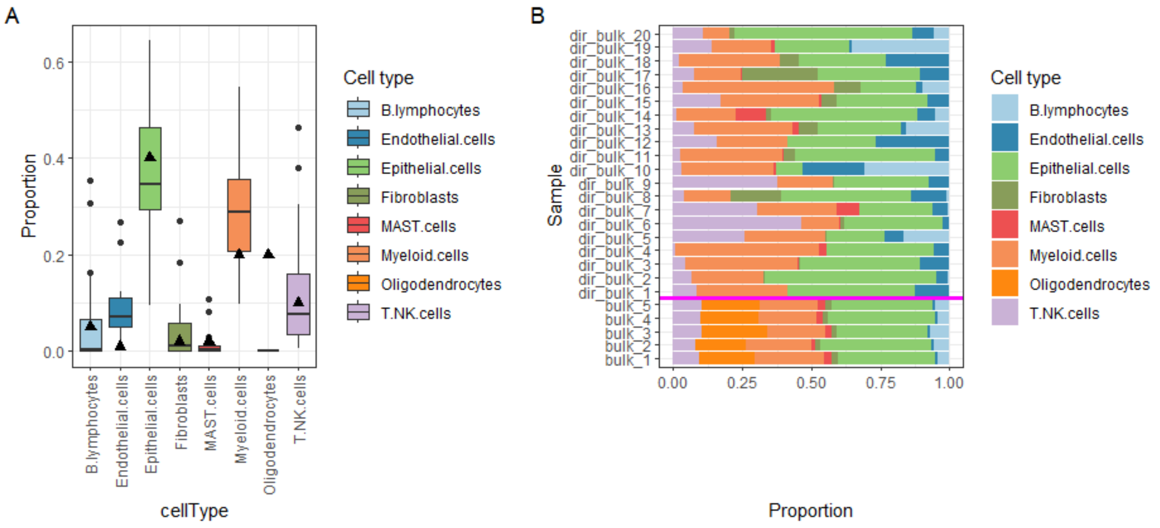


**Figure S2 The setting for one unknown cell type (oligodendrocytes) in simulation study 2.**

(A) The boxplot of Dirichlet distribution-based random cell type proportions. They are subsequently utilized in generating Dirichlet-pseudo-bulk RNA-seq data. The black triangle denotes the true proportion of each cell type, including the unknown cell type oligodendrocytes. (B) The bar plot of two groups of simulated bulk RNA-seq samples, including 5 pseudo-bulk RNA-seq samples, each with 6 cell types (located at the bottom of the bright pink line and used to mimic real bulk RNA-seq) and 20 random-pseudo-bulk RNA-seq samples, each with 5 cell types (located at the top of the bright pink line and used for marker gene mining on unknown cell type). It is obvious that the one cell type in orange is missed in the pseudo samples.

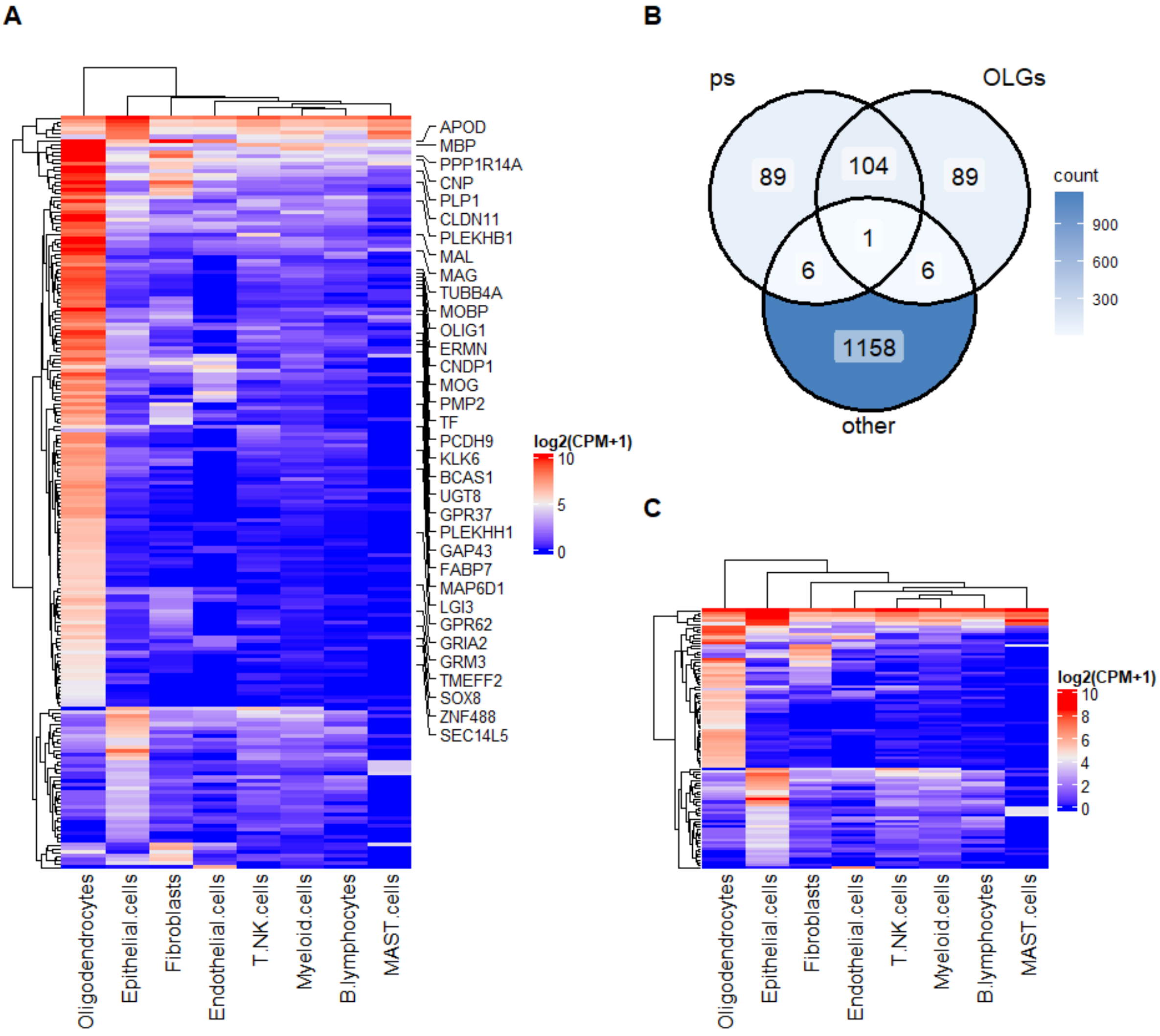


**Figure S3**. Identification of the pseudo marker genes for one unknown cell type (oligodendrocytes) in simulation study 2. (A) Heatmap of psMarker gene expression using the scRNA-seq data on metastatic brain tumor samples from metastatic lung adenocarcinoma (referred to as complete scRNA-seq data). Gene names are shown for genes that are known as marker genes of oligodendrocytes based on the public cell type annotation database. In this simulation study, we know the unknown cell type is oligodendrocytes, and the matched and complete scRNA-seq data is obtainable. (B) The Venn diagram of three gene sets, ps (psMarker genes of unknown cell types identified by MiCBuS via scRNA-seq data from primary lung tumor samples and metastatic bulk data), and scMarker (OLGs: oligodendrocyte marker gene; other:

marker gene of the other seven known cell type identified by scRNA-seq data from metastatic brain tumor samples.) (C) Heatmap of psMarker (unique 95 genes from Venn diagram in (B)) gene expression using the scRNA-seq data from metastatic brain tumor samples.

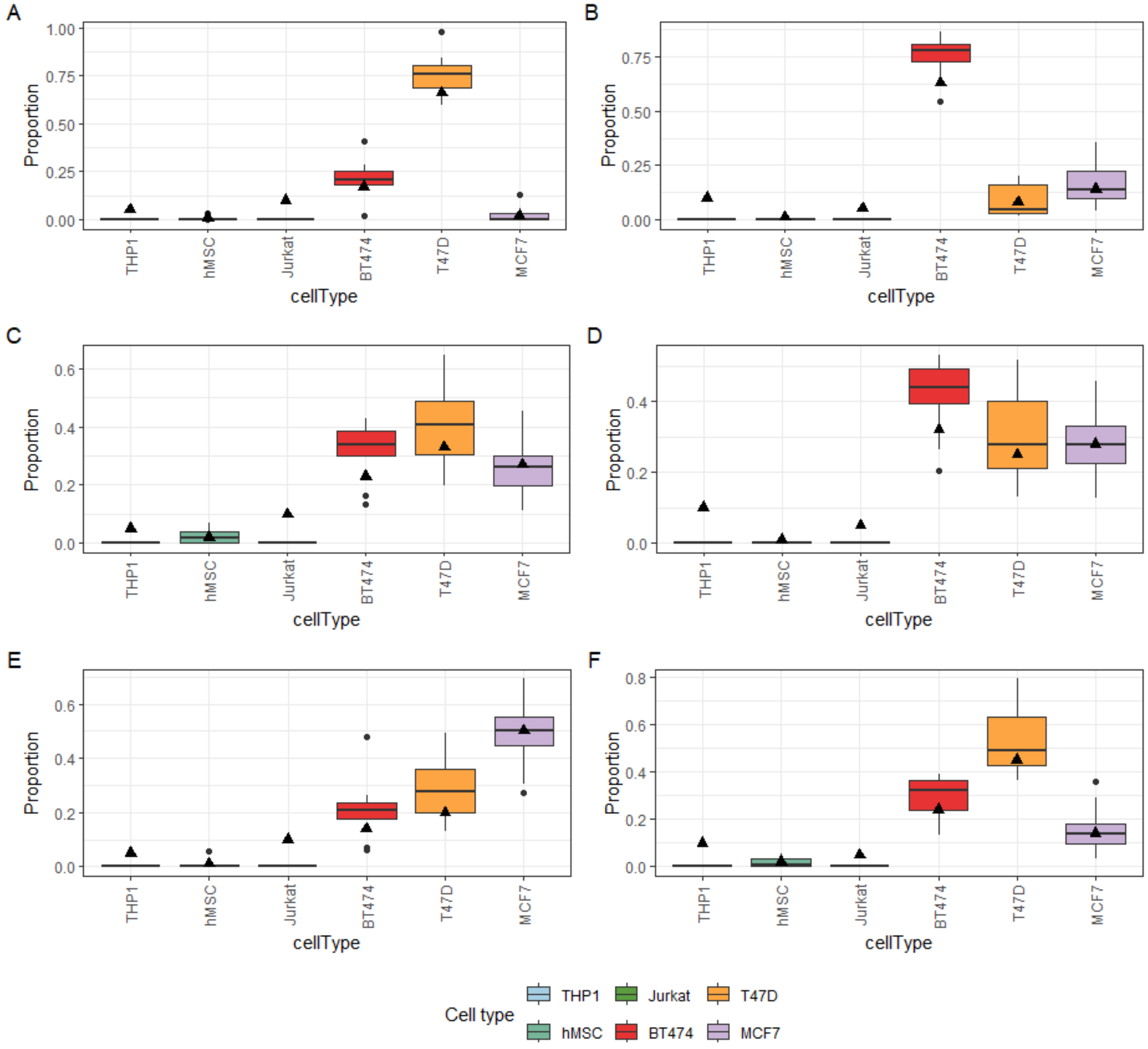


**Figure S4** The boxplot of Dirichlet distribution-based random cell type proportions in setting 2 of real data analysis 1 (THP1 and Jurkat cells unknown). They are subsequently utilized in generating Dirichlet-pseudo-bulk RNA-seq data. The black triangle denotes the golden true proportion of each cell type by cell counting, including the unknown cell types THP1 and Jurkat cells. The data include six different types of mixture, each containing 3 replicates. (A) to (F) correspond for mixture 1 to mixture 6.

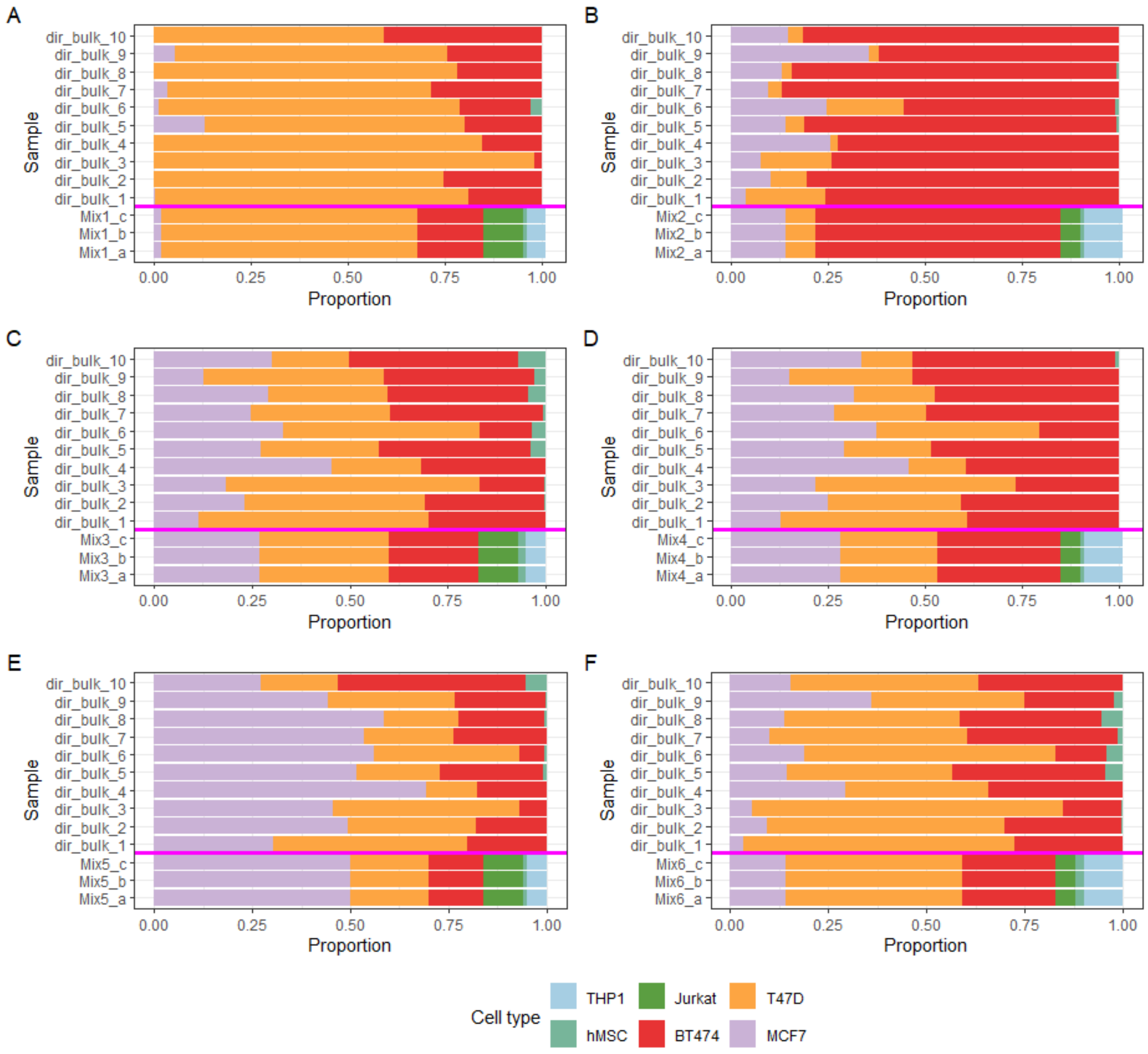


**Figure S5** The bar plot of two groups of bulk RNA-seq samples in setting 2 of real data analysis 1 (THP1 and Jurkat cells unknown). Each panel includes 3 real original bulk RNA-seq samples, each with 6 cell types (located at the bottom of the bright pink line) and 10 Dirichlet-pseudo-bulk RNA-seq samples, each with 5 cell types (located at the top of the bright pink line and used for marker gene mining on unknown cell type). It is obvious that the cell types in blue and dark green are missed in the pseudo samples. (A) to (F) correspond for mixture 1 to mixture 6.

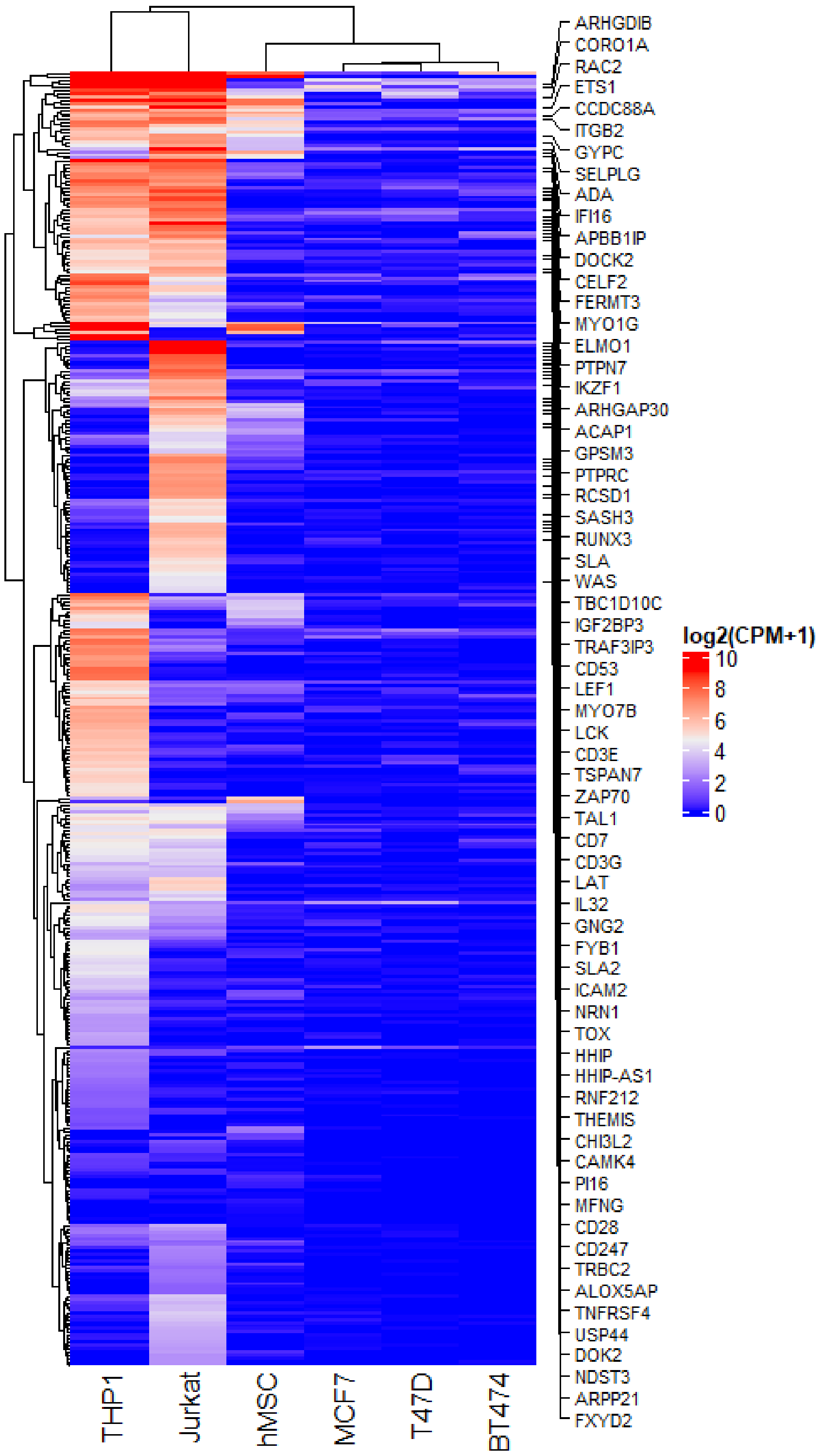


**Figure S6** Identification of the pseudo marker genes in setting 2 of real data analysis 1 (THP1 and Jurkat cells; Supplemental figure for Figure 2.26B). Heatmap of psMarker gene expression using the complete scRNA-seq data. Gene names are shown for genes that are known as marker genes of the Jurkat cell line based on the cell type annotation database. In this setting, the masked unknown cell types are THP1 and Jurkat cells.

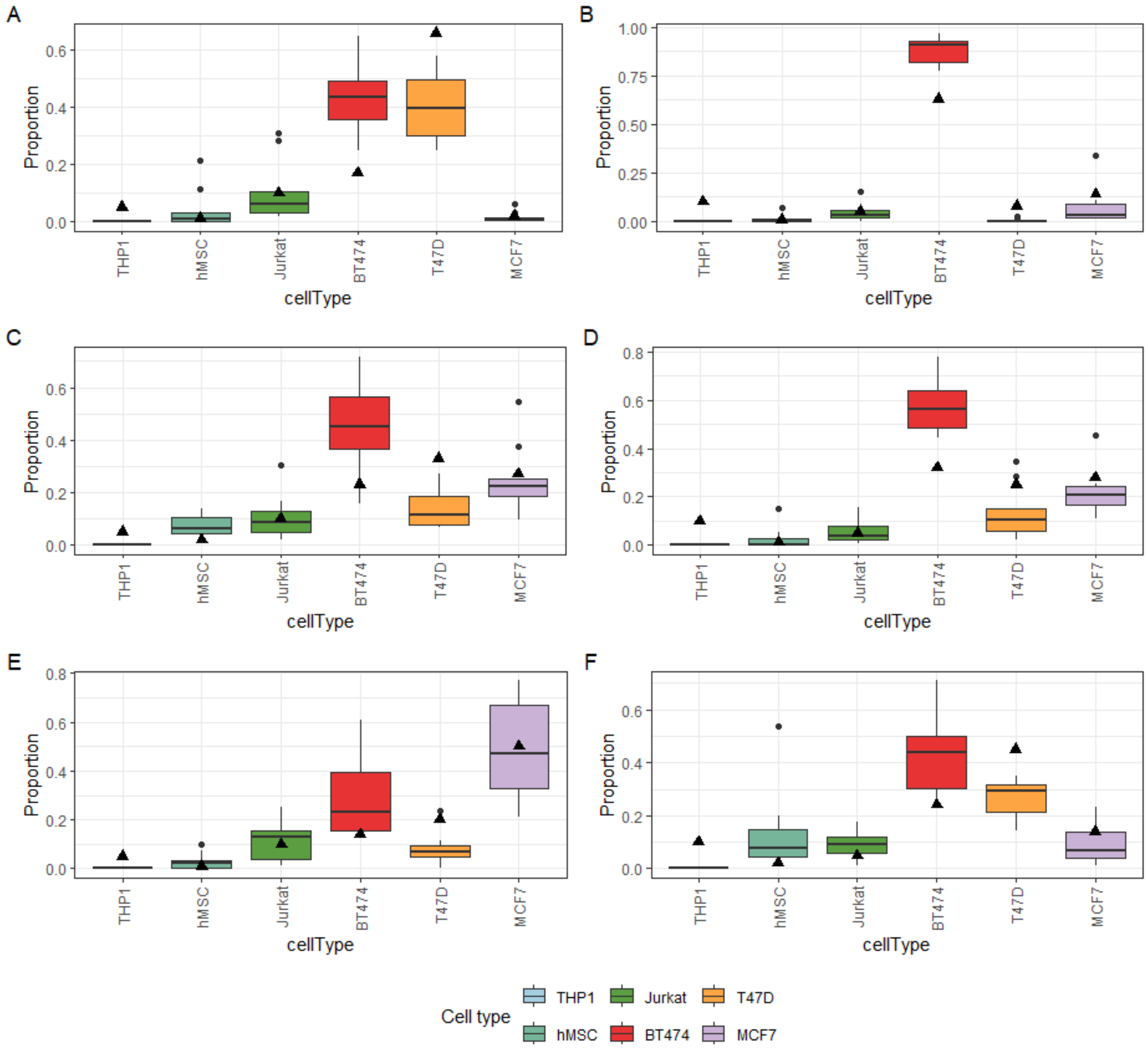


**Figure S7** The boxplot of Dirichlet distribution-based random cell type proportions in real data analysis 2 (THP1 cells unknown). They are subsequently utilized in generating Dirichlet-pseudo-bulk RNA-seq data. The black triangle denotes the golden true proportion of each cell type by cell counting, including the unknown cell type THP1 cells. The data include six different types of mixture, each containing 3 replicates. (A) to (F) are corresponding for mixture 1 to mixture 6.

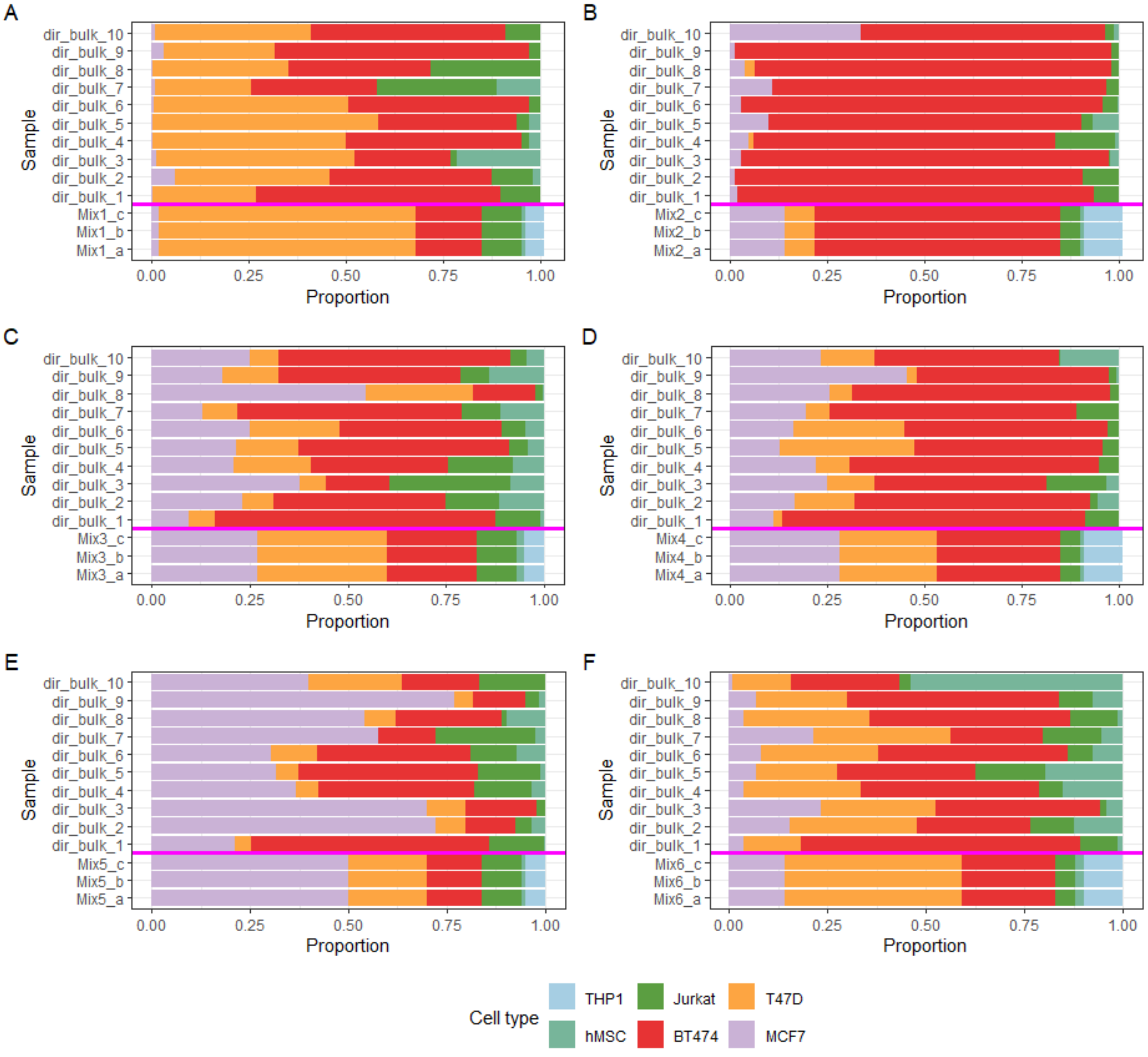


**Figure S8** The bar plot of two groups of bulk RNA-seq samples in real data analysis 2 (TPH1 cells unknown). Each panel includes 3 real original bulk RNA-seq samples, each with 6 cell types (located at the bottom of the bright pink line) and 10 Dirichlet-pseudo-bulk RNA-seq samples, each with 5 cell types (located at the top of the bright pink line and used for marker gene mining on unknown cell type). It is obvious that the one cell type in blue is missed in the pseudo samples. (A) to (F) correspond for mixture 1 to mixture 6.

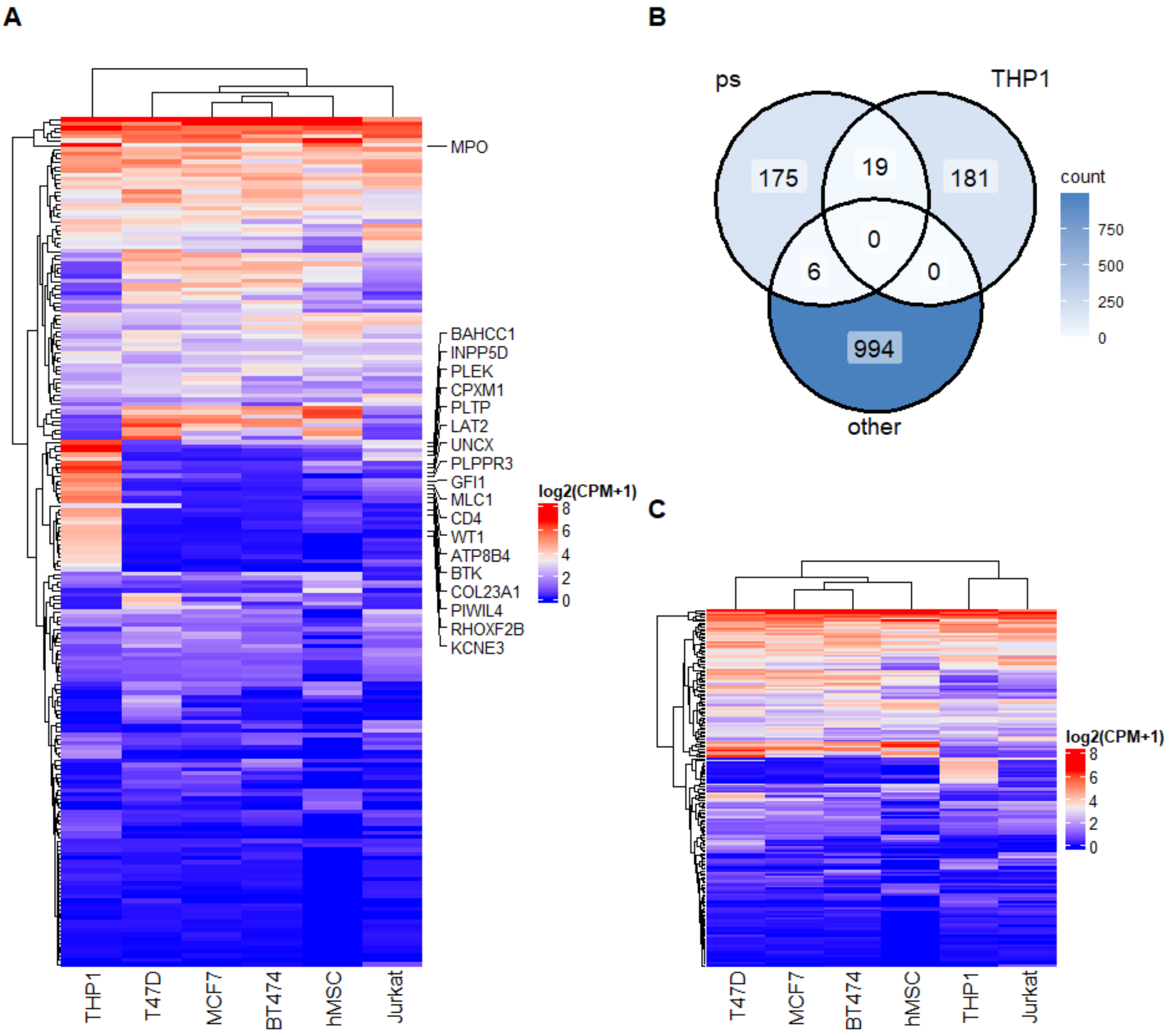


**Figure S9 Identification of the pseudo marker genes in real data analysis 2 (THP1 cells unknown).**

(A) Heatmap of psMarker gene expression using the complete scRNA-seq data. Gene names are shown for genes that are known as marker genes of THP1 cells based on the cell type annotation database. In this setting, the masked unknown cell type is

THP1 cells. (B) The Venn diagram of three gene sets, ps (psMarker genes of unknown cell types identified by MiCBuS), and scMarker (THP1: THP1 marker gene; other: marker gene of the other known cell type identified by complete scRNA-seq data.) (C) Heatmap of psMarker (unique 181 genes from Venn diagram in (B)) gene expression using the complete scRNA-seq data.

**References**

Avila Cobos, F., *et al.* Benchmarking of cell type deconvolution pipelines for transcriptomics data. *Nat Commun* 2020;11(1):5650.

Baron, M., *et al.* A Single-Cell Transcriptomic Map of the Human and Mouse Pancreas Reveals Inter- and Intra-cell Population Structure. *Cell Syst* 2016;3(4):346-360 e344.

Cobos, F.A., *et al.* Effective methods for bulk RNA-seq deconvolution using scnRNA-seq transcriptomes. *Genome Biol* 2023;24(1):177.

Dong, M., *et al.* SCDC: bulk gene expression deconvolution by multiple single-cell RNA sequencing references. *Brief Bioinform* 2021;22(1):416-427.

Kim, N., *et al.* Single-cell RNA sequencing demonstrates the molecular and cellular reprogramming of metastatic lung adenocarcinoma. *Nat Commun* 2020;11(1):2285.

Lu, Y., Chen, Q.M. and An, L. Semi-reference based cell type deconvolution with application to human metastatic cancers. *NAR Genom Bioinform* 2023;5(4):lqad109.
